# Evidence for post-allopolyploidy genetic exchanges between duplicated regions in three ancient polyploidies

**DOI:** 10.64898/2026.06.20.733495

**Authors:** Amrit Kaur Dhillon, Hasini Pasagadugula, Isabelle Pitts, Manav Rohilla, Gavin C. Conant

## Abstract

Many successful lineages, including flowering plants and vertebrates, owe some of their evolutionary prosperity to whole genome duplications (WGD). However, in the immediate aftermath of a WGD, the new polyploid species that is formed often experiences multivalent pairings during meiosis, which can produce inviable gametes. To mitigate the potential harm caused by such pairings, most lineages eventually undergo “diploidization” to restore typical bivalent pairing. A key component of this process is the loss of duplicated genes. While diploidization was once thought to be rapid, recent analyses of polyploidies suggest the process may be more drawn out, with multivalent pairing persisting long after the initial WGD event. Here, we assess evidence for “late” diploidization after three different polyploidies: the teleost genome duplication (TGD), nested polyploidies in *Paramecium* lineages, and the ancient WGD in baker’s yeast. Using our tool POInT (the Polyploidy Orthology Inference Tool), we model the resolution of these events. By analyzing discordance between expected species trees and observed gene trees, we argue that late diploidization was a likely feature in the resolution of all three polyploidies.

## Introduction

Many taxonomically diverse eukaryotic lineages descend from species first formed through polyploidy events, which are also known as whole-genome duplications (WGD). Examples include all the vertebrates (with several additional events among teleost fishes), as well as yeasts, ciliates, and, most strikingly, the flowering plants (1). A new polyploid species can form through the merging of genomes either from the same species or from differing species. In the first case, we refer to the polyploid as an *autopolyploid*, while the second is known as an *allopolyploid* (2–4). Particularly in the case of allopolyploids, we can distinguish two subgenomes that coexist in an extant organism but that are derived from two distinct progenitor species.

Polyploidy can present problems in the production of gametes since there are now four similar copies of every chromosome in each nucleus rather than two. This redundancy introduces the potential for cross-over events between various combinations of those chromosomes (5–7), so the resulting chromosome associations are referred to as being *tetravalent* or *multivalent* rather than the bivalent pairing seen in typical diploid taxa. Multivalent pairing is more common in new autoploids than new alloploids due to the presence of four identical copies of the chromosomes. However, there exist many examples of new autopolyploids that instead show conventional bivalent pairing and many allopolyploids with some multivalent pairing (6). Multivalent pairings can result in various deleterious outcomes involving the formation of inviable gametes (8). As a result, lineages arising from ancient polyploidies (paleopolyploids) will have resolved their multivalent pairings (6,9,10) and are referred to as having *diploidized* because they show almost exclusively bivalent pairing.

When the period prior to diploidization in allopolyploids includes multivalent pairings, those pairings can drive complex events in chromosome evolution. For instance, these pairings can result in the replacement of an entire chromosome from one of the contributing subgenomes with the corresponding chromosome from the other subgenome (11). A replacement of this type will result in that set of four chromosomes appearing to be due to autopolyploidy rather than an allopolyploidy. Ancestral allopolyploids that have undergone some of these homoeologous replacements can then be referred to as *segmental allopolyploids* (12).

Curiously, a few polyploid lineages do not seem to follow the diploidization pattern just described. The most well-known of these taxa are the salmonids, which are derived from an ancient tetraploidy. In these fishes, multivalent (*tetravalent* in this case) pairings are currently ongoing in some regions of certain chromosomes (13,14). There are several interesting features related to these pairing patterns. For instance, the polyploidy-produced duplicates (known as *ohnologs*) (17) in tetravalent regions show less sequence divergence than do surviving ohnologs in other regions (15). Moreover, careful comparisons of gene trees to the expected species tree in the genomes sharing this event suggests that the loss of tetravalent pairing occurred at different times and in different chromosomal regions in each lineage (16). Finally, the continued pairing of homoeologous chromosomes suppresses duplicate losses, resulting in more surviving ohnologs in those chromosomal regions. A knock-on effect of this phenomenon is that gene conversion events between these ohnologs keep their sequences very similar within a species (18).

In recent work, we have suggested that incomplete diploidization also characterizes several other ancient polyploids, namely those in carp, in sturgeon and in apples and pears (18). In other words, genomes in these lineages also continue to experience some degree of homoeologous recombination, presumably through multivalent pairings. Our conclusions were consistent with previous analyses regarding the distribution of the observed gene trees and divergences seen in some of these lineages (9,10,19).

These results raise the question of whether these fishes and the apples and pears are in fact unique in experiencing homoeologous exchanges long after the polyploidy event itself or if this pattern of long-lived homoeologous exchanges is common. This second alternative could well be masked in extant taxa if the exchanges ceased at some point in the past. And in fact, there is some evidence that homoeologous exchanges may have persisted for a significant period after other polyploidies: analyses of the gene trees from extant teleost fishes suggest that this lineage may have experienced reasonably frequent homoeologous exchanges early in its history (20). Here, we assess several lines of evidence for continued recombination events after three ancient polyploidies: the teleost genome duplication (TGD) (21,22), the duplication in bakers’ yeast and relatives (23), and that in several ciliate species (24). We find suggestive indications of such events using different lines of evidence, although that evidence differs noticeably from event to event.

## Results

### Analysis of three ancient polyploidies using POInT

We generated blocks of double-conserved synteny (DCS) for ten to eleven genomes sharing three separate tetraploidy events, one in yeasts, one in bony fishes, and one in the ciliates (*Methods*). In each group, an ancestral gene duplicated by WGD and now represented by at least one homolog of that gene in each extant species is referred to as a *pillar*. For the TGD (Figure 1A), we expanded our previous dataset from 8 species and 5,589 pillars (25) to 11 species and 7,669 pillars. For paramecia, our analysis encompasses 10 species and 3,925 pillars, verses 3 species and 11,683 previously (26). For the ciliates, we separated the ancient and recent WGD events with POInT’s octoploid models (Supplemental Figure 1, *Methods*). In both cases, we used an approach that minimized the number of synteny breaks across the extant genomes in these DCS blocks to infer an approximate ancestral gene order that existed just prior to the WGD events (*Methods*). These orders were used for our POInT analyses and for analyses of the distribution of gene trees in the polyploid taxa (see below). We have previously described both the 11 species and the inferred ancestral order used in the yeast dataset (27,63).

**Figure 1:**
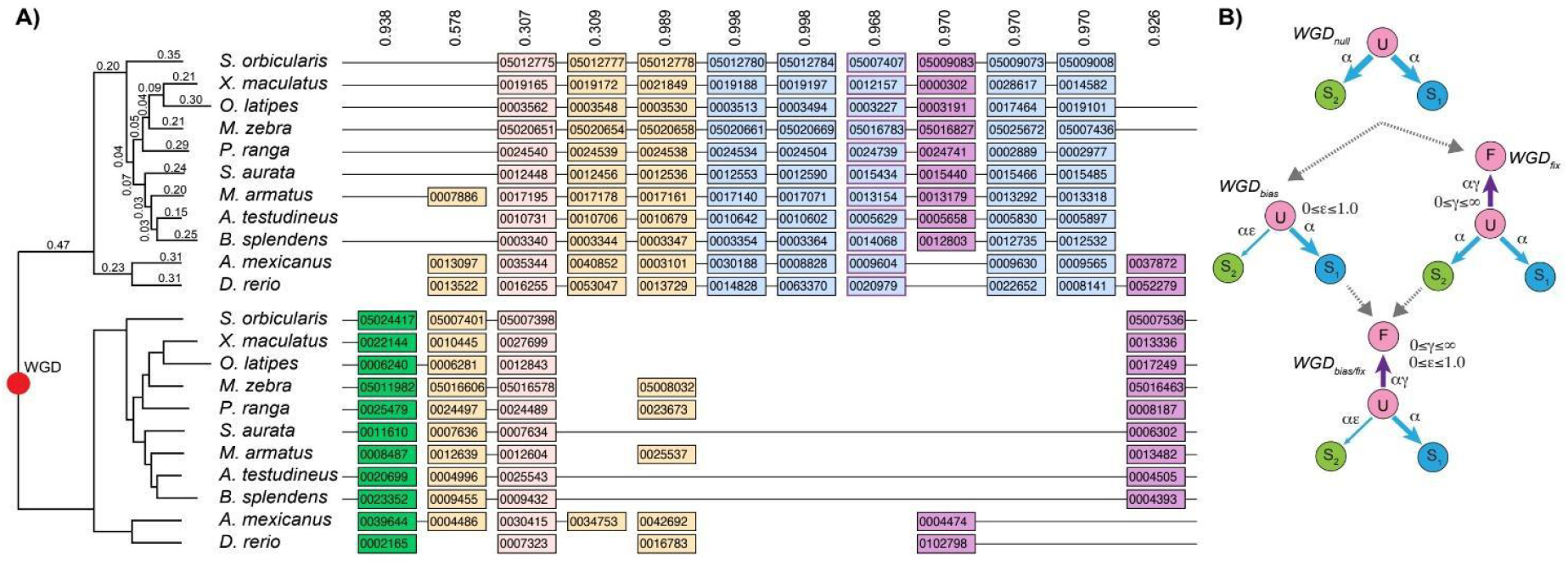
**A)** Regions of DCS (double-conserved synteny) in eleven genomes sharing the ancient teleost genome duplication (TGD). The two mirrored species trees at left show our best inference of the relationships of these species and their respective genes duplicated at this event. There are 2^11^=2048 possible organizations of these genomic regions into orthologous chromosomal regions; POInT computes the likelihood of all these possibilities and depicts the most probable. The numbers at the top of each pillar represents the proportion of the total probability accounted for by the depicted orthology relationship; hence numbers close to one indicate regions of high confidence in the depicted orthology. Given that orthology inference, we can consider whether the ohnologs created by the TGD survive. In many cases, some have been lost and others retained (tan pillars). In a few cases, all species retain the duplication (light pink pillar). If all species have lost one gene copy, there are three possible patterns that might be observed. If all copies were lost from the less-favored subgenome, we represent that in blue: if all copies were lost from the favored subgenome, that possibility is shown in green. Finally, if some species lost the genes from the favored subgenome and some from the less-favored subgenome, that *reciprocal gene loss* is shown in magenta. This image can be regenerated at https://wgd.statgen.ncsu.edu. **B)** Four models of ohnolog loss after polyploidy. In the WGD_null_ model at the top, ohnologs are lost at an equal rate from both ancestral subgenomes. Allowing ohnologs to become fixed at a relative rate γ generates the WGD_fix_ model at the right, while allowing the model to estimate the favored subgenome from the loss data generates the WGD_bias_ model, where losses from that favored subgenome occur at a reduced rate ε (0□ε□1). Combining these two models produces WGD_bias/fix_ with both processes (bottom). These models can be compared with a likelihood ratio test (*Methods*).

We analyzed these DCS datasets using nested models of duplicate loss and fixation (Figure 1B) using POInT (v. 1.63). For the TGD, the data suggested the occurrence of both duplicate pair (ohnolog) fixation and biases in subgenome losses (*P<*10^−20^; Figure 1B). These biases mean that the subgenomes in this event are evolutionarily distinguishable, and hence that the event was an allopolyploidy. For the paramecium WGD, the evidence for fixation was very strong (WGD_fix_ vs. WGD_null_, WGD_bias/fix_ vs. WGD_bias_, *P<*10^−20^ in both cases). However, the evidence for bias was less strong (WGD_bias_ vs. WGD_null_, WGD_bias/fix_ vs. WGD_fix_, *P<*10^−4^ in both cases). The level of bias was also inferred to be relatively weak (ε=0.86, meaning about a 15% advantage for subgenome #1). As a result, we have chosen to use the WGD_fix_ model in our analyses of this event. In previous work, we also showed that there was little evidence for bias in subgenome losses after the yeast WGD (27), but fixation was still strongly detected (*P<*10^−10^).

### Species trees inference with POInT

Although published topologies are available for the species analyzed (28–30), we sought to explore some of the topological space of species trees under the POInT gene loss models. We hence inferred gene trees for all single-copy genes in the three datasets and ranked them by their topological frequency (*Methods*). We then assessed how well each of the most-frequent topologies fit the synteny/gene loss data using POInT. Figure 2 shows the topologies tested for each event and the one with the highest likelihood of having generated the loss patterns seen (i.e., the ML tree under the POInT models).

**Figure 2:**
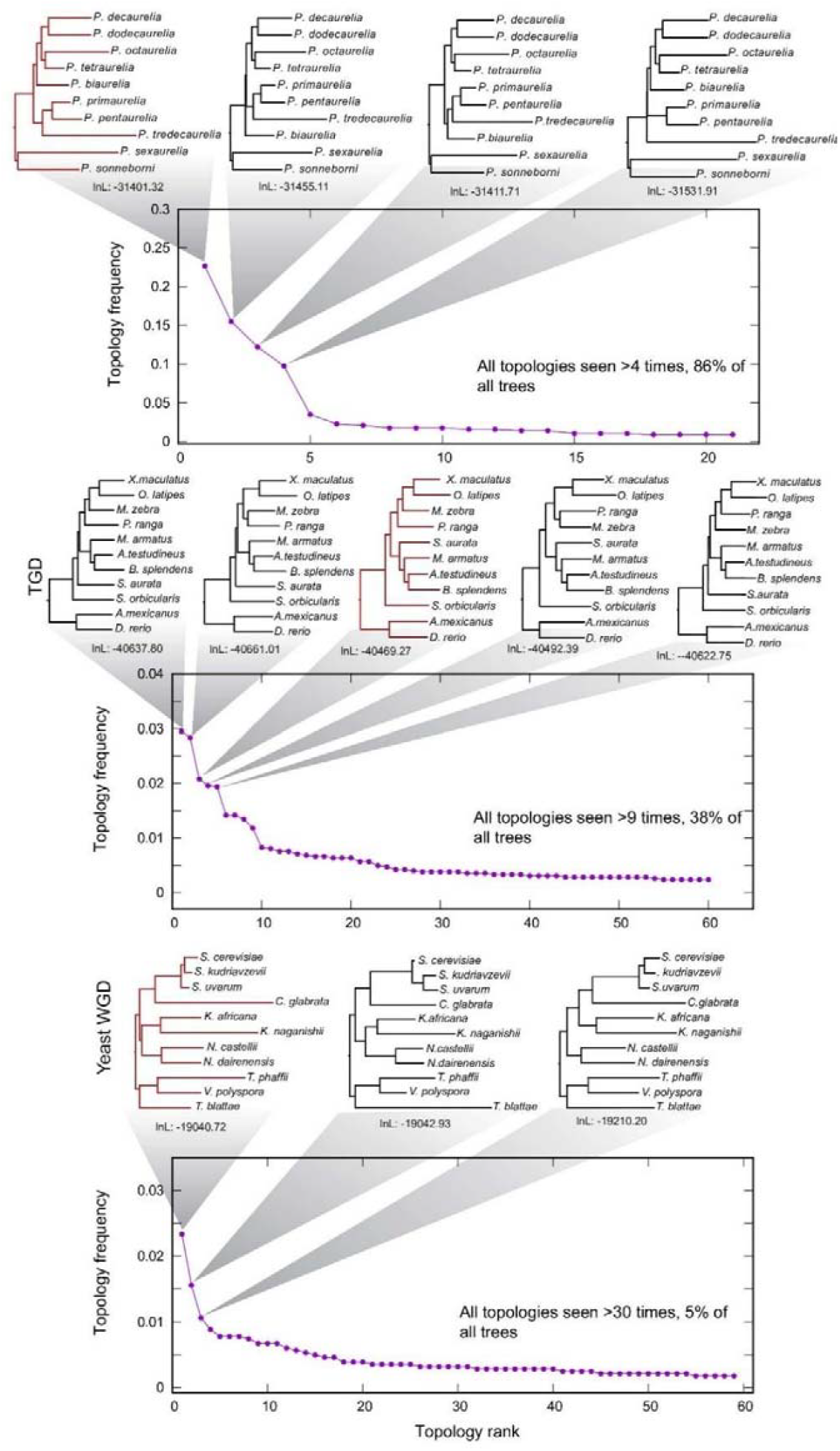
The distribution of observed gene tree topologies for the three events. We ranked all the observed single-copy gene tree topologies (*x*-axis) in the order of the frequency with which different pillars supported them (*y*-axis). We then fit the most common topologies to the DCS data using POInT. The resulting POInT likelihood values (and associated trees) are shown above each plot, with the maximum likelihood topology shown in red. This topology was used for all remaining analyses. **A)** Paramecium WGD, **B)** Teleost genome duplication (TGD), **C)** Yeast genome duplication.

### Ohnolog losses were spatially clustered over the history of these polyploidies

POInT models the resolution of a WGD with the set of model states shown in Figure 1B. For each pillar in each event, we computed the conditional probability of every possible state transition change along every branch of the species tree (*Methods*). These state-change values allow us to assess the probability that an ohnolog loss occurred at each pillar on any given branch of the tree. We then compared the observed loss probabilities to what was seen when the order of the pillars was randomized. For all the early branches of the trees (and most of the later ones), there was significantly more similarity in the loss patterns of adjacent pillars than would be expected by chance (*P<*0.05, *Methods*, Figure 3). These results could have been driven by one or both of two processes. First, ohnolog losses could have occurred in blocks of genes (31). Second, homoeologous recombinations could have ceased later in time in some regions of each chromosome than in others. In this second case, places where recombination ceased later would have had less subsequent time for ohnolog losses, giving rise to regions with more surviving ohnologs.

**Figure 3:**
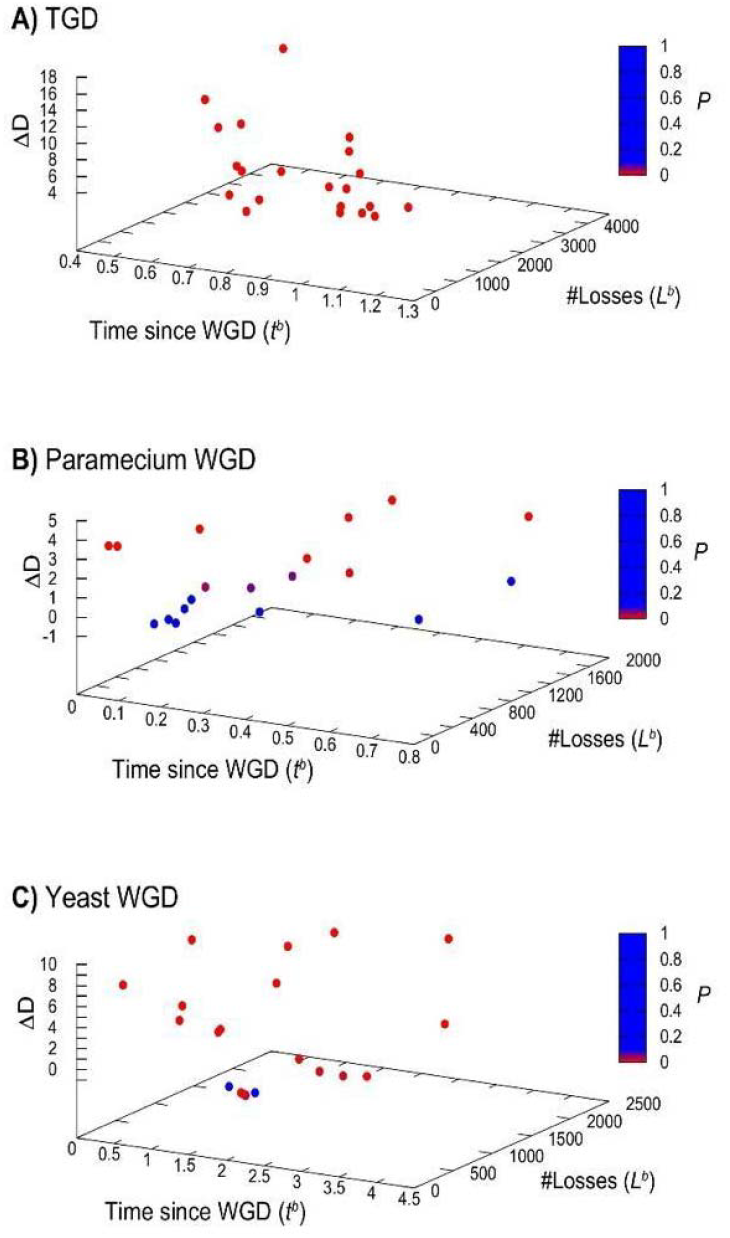
Many phylogenetic branches show excess clustering of ohnolog losses above what would be expected by chance. On the *x*-axis is the sum of the branch lengths from the branch in question back to the root (corresponding to the total model time since the WGD); on the *y*-axis is the number of losses experienced along that branch (*L;*equation 6). On *z* is the degree of difference between the observed similarity in losses between neighboring pillars and that expected by chance (Δ*D*; equation 7; *Methods*). Points are color-coded by the significance of the differences between the observed similarity and that seen in the randomized pillars (color code at right).

### Gene trees are non-uniformly distributed along the ancestral chromosomes for the TGD but not for the paramecium and yeast WGDs

We computed the gene tree for each pillar in our datasets *(Methods*). After excluding pillars with low orthology confidence *c*(0 ≤ *c* < 0.90), we compared these gene tree topologies to the expected species tree using the Robinson-Foulds (RF) distance (32). We then asked if the resulting distances between these gene trees and the species (*RF*_*i,spp*_) were uniformly distributed along the ancestral genome order (*Methods*). To do so, we computed the absolute value of distance difference between neighboring pillars (*RF*_*i,spp*_ − *RF*_*i,spp*_) and computed the mean of those differences:

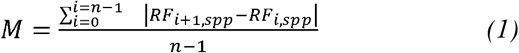

We then randomized the pillars 1000 times and computed *M* for these randomized pillars to assess whether there was local structure in the variation in gene tree topologies along the ancestral order. For the TGD, we found that neighboring pillars showed more similar distances to the species tree than would be expected by chance (*P*< 10−3) but saw no such effect for the paramecium and yeast WGDs (*P*> 0.05, Figure 4A-C).

**Figure 4:**
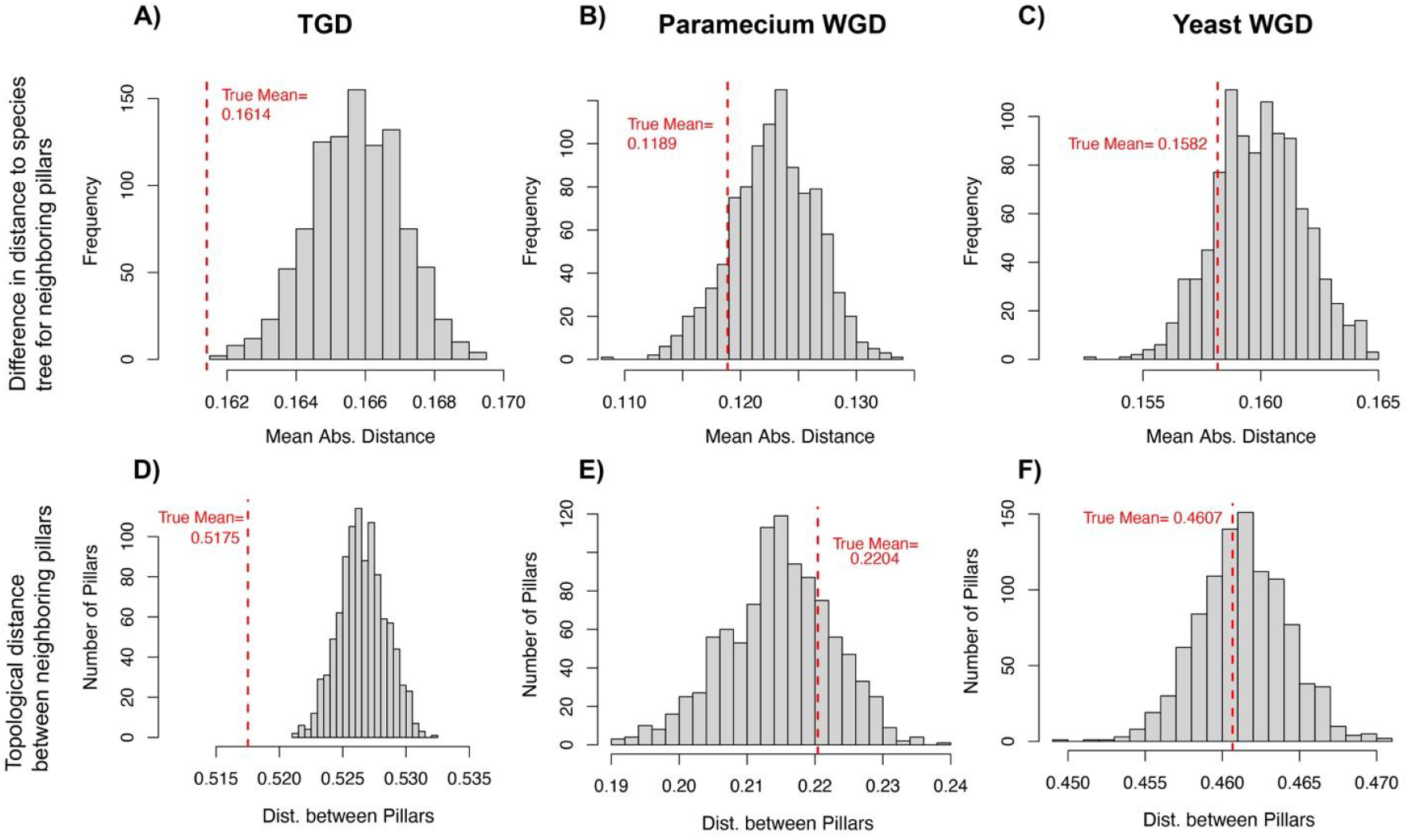
**(A-C)** Distribution of distances to the species tree observed for gene tree topologies: For each polyploidy event, the (mean absolute distance) *M* between the gene trees and the species tree was computed (Equation 1). This value is shown as the vertical dashed red line. It is compared to the distribution of 1000 random means produced through pillar randomization (*Methods*), shown as histograms. All the true means lie to the left of the simulated means, but only TGD shows a value outside of those seen in simulation (*P* < 10^−3^). **A)** TGD, **B)** Paramecium WGD, **C)** Yeast WGD. **(D-F)** Topological distances between neighboring gene trees. Here *M*_*n*_ is the mean distance between pairs of neighboring single-copy gene trees, shown as a vertical dashed line. Histograms again show the distribution of means seen after pillar randomization (*Methods*). Again, the actual mean seen for the TGD does not fall within the range of values seen in simulation (*P* < 10^−3^). **D)** TGD, **E)** Paramecium WGD, **F)** Yeast WGD.

We next asked whether neighboring, single-copy, genes showed similar topologies to each other, regardless of their relationship to the assumed species tree. We computed the mean RF distance (*M*_*n*_) between pairs of neighboring pillars:

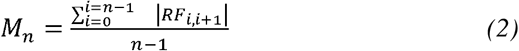

We again compared *M*_*n*_ to that seen after 1000 randomizations of the pillar order, and once again, the TGD showed significantly smaller distances between neighboring pillars (*P*< 10^−3^), but the paramecium and yeast WGDs did not (*P* > 0.05, Figure 4D-F).

### Suggestion of early gene conversions after ancient allopolyploidies

If homoeologous recombinations persisted for some time after the respective allopolyploidies, we might expect that some of the gene trees inferred might preserve a history of gene conversion events associated with those recombinations. In other words, the last common ancestral event for some of the ohnolog pairs may not have been the allopolyploidy itself, but rather a more recent homoeologous recombination. To test this possibility, we proposed three possible tree topologies that retain duplicates from the allopolyploidy (Supplemental Figure 2). The first topology consists of a pair of mirrored species trees consistent with no post-WGD recombination events. The second topology posits universal recent gene conversions, such that each member of an ohnolog pair is sister to its own paralogous copy from the same genome (Supplemental Figure 2). In other words, every ohnolog pair has undergone a complete gene conversion with each other since the most recent speciation event. Finally, the third topology hypothesizes a gene conversion event having occurred in one or both of the two clades produced by the first post-WGD split in the species tree. In other words, after the first observed speciation event following the WGD, the common ancestor of all the lineages on one or both sides of that split experienced a recombination event. The result of that event or events will be that both of those lineages will have ohnologs that cluster with each other in the unrooted gene tree, giving the erroneous impression of two independent WGD events at that pillar (Supplemental Figure 2).

We drew an inference of gene conversion if the inferred gene tree was closer in R-F distance to one of the gene conversion topologies than it was to the topology with mirrored species trees. To assess the statistical significance of this conclusion, we made 100 simulations of sequence evolution from the species tree for that pillar. If those simulations never produced an improvement in likelihood for the gene conversion topology compared to the species tree as large as was seen in the real data (*Methods*), we concluded that the support for the hypothesis of gene conversion was significant. For the TGD, a relatively small proportion of pillars showed such evidence, but for the yeast and paramecium events, we observed a relatively high proportion of pillars with evidence for early gene conversion events (Table 1). Permutation tests also suggested that these events were clustered in the ancestral genome order (*Methods*, Table 1).

**Table 1:** **a)** Total cases where one of the GC trees fit the aligned sequences better than did the species tree (*Methods*) out of the total number of pillars where the three trees were distinct and could be compared. In parentheses is the breakdown between the early GC and the full GC cases. We required an improvement in log-likelihood of 5 units for the GC tree for these cases. **b)** As in the previous column, but with a cutoff of at least 10 log-likelihood units rather than 5. **c)** For the likelihood cutoff of 5, the probability that randomly selected pillars being on average closer to each other than observed for these GC events (*Methods*).

| Event | Gene conversions/Total Pillars<br>(Early/Full)<sup>a</sup> | Gene conversions with<br>$\Delta\ln L \geq 10$ (Early/Full)<sup>b</sup> | P-value GC<br>clustering<sup>c</sup> |
| --- | --- | --- | --- |
| TGD | 39/1100 (31/8) | 34 (26/8) | 0.57 |
| Yeast WGD | 56/311 (56/0) | 38 (38/0) | 0.02 |
| Paramecium WGD | 125/327 (122/3) | 120 (117/3) | <0.01 |

### Pairwise Estimates of Gene Conversion

For each of the polyploidy events, we also assessed the evidence for very recent gene conversion events by comparing each ohnolog pair to the ortholog of one of those genes in a near relative species (Figure 5; *Methods*). The result is a triplet of sequences with two ohnologs and the ortholog of one (inferred from the POInT synteny data). As shown in Figure 6, in the absence of gene conversion, the focal gene should have a sequence closer (in terms of nonsynonymous divergence or *K*_*a*_) to its ortholog than to its paralog/ohnolog. The reason is that the speciation event is much more recent than the ancient WGD event. Cases where the focal gene is significantly closer to its paralog from the WGD can be attributed to recent gene conversion (*Methods*). As shown for selected examples in Figure 5, putative cases of gene conversion are also associated with low synonymous divergence (*K*_*s*_) between the ohnologs of the pair. (Data for all taxa pairs is given in Supplemental Table 1 and Supplemental Figure 3as a very short root branch in the species as a very short root branch in the species −5).

**Figure 5:**
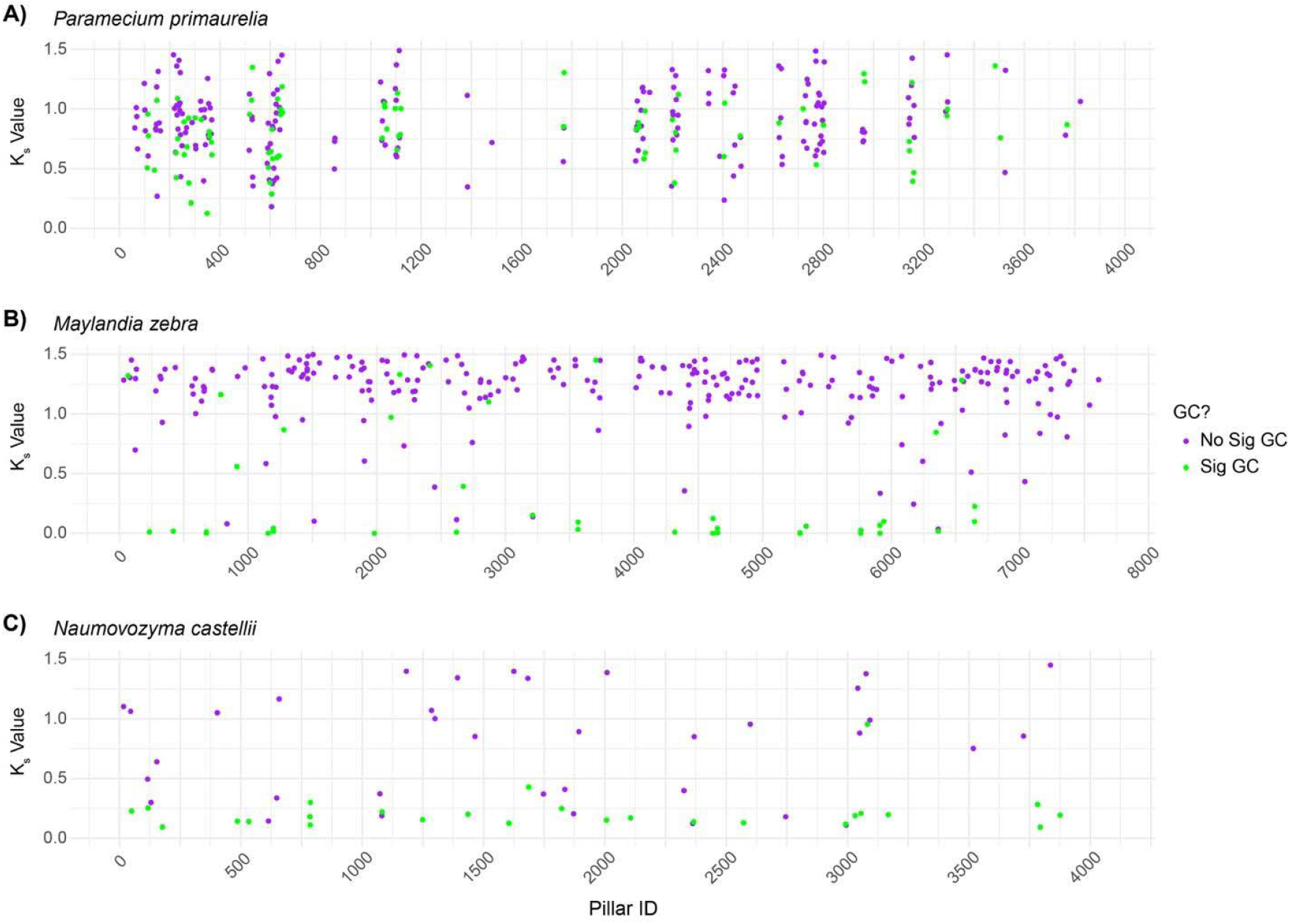
Recent gene conversion in selected taxa from each of the three polyploid events. Statistically significant cases of gene conversion are indicated in green (*P* < 0.05); all other ohnolog pairs are shown in purple. The *x* axis shows the position in ancestral genome order; on *y* is the synonymous divergence between the ohnolog pair (*K*_*s*_, or the number of synonymous substitutions per synonymous site, *Methods*); ohnolog pairs with K_s_ greater than 1.5 are omitted. Each plot displays pillars of ohnologs from POInT with high confidence (90%) orthologs (Equation 8, *Methods*). **A)** *Paramecium primaurelia* ohnolog pairs tested against *Paramecium tredecaurelia* as the outgroup (101 / 3925 pillars had GC ohnologs with high confidence orthologs). **B)** *Maylandia zebra* ohnolog pairs tested against *Xiphophorus maculatus* as the outgroup. (60 / 7669 pillars had GC ohnologs). **C)** *Naumovozyma castellii* ohnolog pairs tested against *Naumovozyma dairenensis* as the outgroup. (104 / 4065 pillars had GC ohnologs).

**Figure 6:**
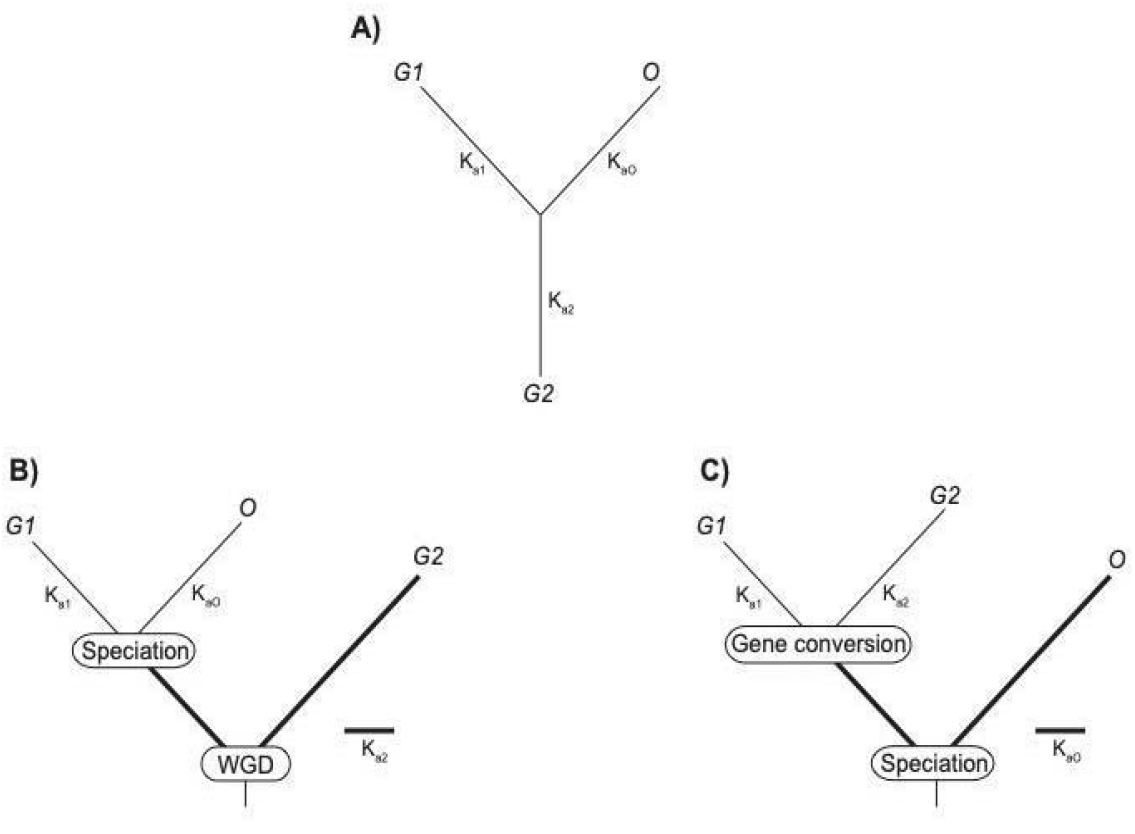
Triplet-based sequence test for gene conversion. **A)** Degenerate phylogenetic tree depicting the time-reversible relationship between the genes *G*1, *G*2, and *O. G*1 and *G*2 are duplicates (ohnologs) of each other and gene *O* represents the ortholog of *G1*. The estimated nonsynonymous divergence (K_a_) that has occurred along each branch is also illustrated. **B)** In the absence of gene conversion, the tree from *A* will collapse to a species tree where the divergence of *G2* is greater than either *G1* or *O*. In other works, *Ka*_2_ is larger than *Ka*_1_ and *Ka*_0_ because *Ka*_2_ represents both its own divergence from the common ancestor and the shared divergence of genes *G*1 and *O* after the WGD and prior to the speciation of these two lineages. **C)** If *G1* and *G2* have experienced significant recent gene conversion, their mutual divergence may be less than either shows to *O*. In this case we expect *Ka*_0_ to be larger than *Ka*_1_ and *Ka*_2_.

## Discussion

Early analyses of genomes sharing ancient genome duplications suggested that these polyploidy events were followed by the rapid loss of many of the duplicated genes created by them (33,34), leading to the suggestion that polyploidy had been followed by a “lag” phase where proper chromosome pairing was restored through ohnolog loss, after which the polyploid population could start to subdivide by speciation (35). Although our analyses of nine paleopolyploidies also supported this rapid loss model (36), there is now increasing evidence that not all ancient polyploidies have followed this pattern.

The salmonids have long been known as an exception to the rapid loss of both duplicated genes and of homoeologous pairing, since they preserve both tetravalent pairing in some chromosome locations (13,14) and have many surviving duplicates with low sequence divergence (15,18). These two patterns strongly suggest a history of progressive loss of tetravalent pairing down to the present day (16). Similar forces appear to have been at work in the sturgeons (9), to the point that independent polyploidies had been inferred in early work before the potential for post-WGD recombination was included in the analyses (37,38). We recently described a total of four polyploidies, including the salmon and sturgeon events, as well as events in carp and apples and pears, that showed patterns of many surviving duplicates and evidence for recent gene conversions, all suggestive of continued homoeologous exchanges down to the present in these groups (18).

Thus, as the study of post-polyploidy evolution has advanced, nuance has been added to the model of rapid diploidization and duplicate loss. Analyses of the distribution of gene trees for duplicates surviving from the teleost genome duplication suggest that this event too was followed by a period of homoeologous exchanges (20), consistent with studies in modern polyploids (39).

Our analyses here support the premise that post-WGD evolution has been characterized by genetic exchanges between the two parental genomes after not only the TGD but also events in yeasts and paramecia. At a high level, the distribution of duplicate gene losses and retentions is non-uniform along the inferred ancestral chromosome orders (Figure 3), a pattern that could be due either to duplicate losses having encompassed more than one gene at once (31) or the local preservation of duplicates by homoeologous exchanges, or both. For the TGD, we see an additional pattern of a non-uniform distribution of gene trees along the ancestral order. Unlike the duplicate loss clusters, these topological differences are more consistent with homoeologous exchanges after polyploidy. However, even stronger evidence for ancient homoeologous exchanges is found in our comparisons of the observed gene trees to the topologies expected with and without ancient gene conversions. For all three polyploidies, there is a set of loci with statistically significant evidence for gene conversion events that post-dated the first speciation event among the taxa studied (Table 1). This result is supported by the patterns of even more recent gene conversions seen especially in the paramecia (Figure 5). (Although recent gene conversions are seen in yeast, we note that previous work has found that these events are almost exclusively confined to the ribosomal proteins and histones (40,41) and probably are not related to homoeologous pairing in the modern species).

The implications of these more complex patterns of post-WGD evolution are numerous. One interesting one lies in our prior claim that the yeast WGD was followed very quickly by speciation events among the extant taxa (42,43). That argument was based on the very small number of shared duplicate loss events between the modern clade that includes *S. cerevisiae* and the clade including *V. polysporus* (Figure 2). This lack of shared losses is represented in POInT’s models as a very short root branch in the species tree inferred (Supplemental Figure 6). But if homoeologous exchanges preserved duplicates for any extended period of time, any dating arguments based around this lack of shared losses is suspect. In fact, salmonids and sturgeons have similarly short root branch lengths estimated by POInT (18). If we consider the yeast data in Table 1, 56/311 of the pillars we analyzed preferred the hypothesis of early gene conversion to the species tree, implying that at least 18% of the genome was still experiencing homoeologous pairings *after* this first speciation event. Since we cannot by definition detect conversion events prior to this first speciation, it may be that a considerable period followed the WGD prior to the speciation event that we do not see in the loss data. We had also previously found that there were almost as many reciprocal losses (where the *S. cerevisiae* lineage lost copy A and the *V. polysporus* lineage copy B) as there were cases of shared loss (33,36). This conclusion, however, loses some force in the case of rampant gene conversion, as those two copies were likely to have been quite similar if they recently experienced gene conversion. Indeed, such conversions may help resolve the paradox that the yeast WGD event is thought to have been an allopolyploidy (44), but it does not show the pattern of biased subgenome losses that other events do (27,45,46).

The patterns of gene conversion and loss clustering seen here are further evidence that a clear distinction between allopolyploidy and autopolyploidy is likely illusory (12) and that the phrase “genome duplication,” while very useful, hides considerable complexity that we are only now starting to be able to unravel. They also reinforce the great utility of having many genomes sharing a particular paleopolyploidy for analysis, as such taxonomic depth allows the type of fine-grained evolutionary analysis necessary to untangle these trends.

## Methods

### Improved synteny datasets for the teleost and paramecium WGD events

Using recently released genomes, we improved our previously described double-conserved synteny (DCS) dataset for the teleost genome duplication (TGD) and paramecium WGD (25,26). Eleven genomes sharing the TGD were obtained from Ensembl release 107 (47); ten genomes for the paramecia were taken from the Paramecium DB (48), as described by Gout et al., (28). The yeast WGD dataset has been previously described (27,43).

The pipeline for generating DCS datasets has three steps. In the first step, we conduct a homology search between the coding sequences of all the genes from each polyploid genome and the coding sequences from an outgroup genome lacking the polyploidy. The outgroup for the TGD was spotted gar (49) and that for the paramecia polyploidy was *Paramecium caudatum* (50). The homology search was carried out with GenomeHistory 2.1 (51), which uses BLASTP (52) to identify those homologs. We required a homolog pair to have a BLAST E-value of 10^−7^ or less and 55% identity or more at the protein sequence level in the TGD. The respective values for the Paramecium WGD were *E* ≤ 10^−6^ and 65% protein identity.

In the second step, we used *POInT_genome_scaffold* (26) to infer the blocks of conserved synteny between each polyploid genome and the outgroup. This tool uses simulated annealing to place genes from the polyploid genome into homoeology relationships with a single outgroup gene. We increased the running time of the simulated annealing search until the optimality score converged for each genome pair. The ciliate genomes have had two tetraploidies in these genomes since the split with *P. caudatum* (28). Therefore, we looked for blocks of quadruple-conserved synteny rather than the DCS blocks of the TGD (see below; “Deconvoluting a nested polyploidy”)

In the third and final step, we merged the optimal scaffold runs from step two to produce a single set of homoeologous genes across all the polyploid genomes. In both cases, we carried out steps 1 and 2 with more than the final 11 and 10 genomes, omitting genomes with poor scaffold scores from step 2 or omitting a genome to produce a final dataset of no more than 11 genomes (the maximum size that is computationally feasible for POInT). For the TGD, these omitted genomes were those of *Gasterosteus aculeatus, Neogobius melanostomus, Takifugu rubripes, Gadus morhua* and *Cottoperca gobio*. For the paramecium event, they were *Paramecium jenningsi, Paramecium novaurelia* and *Paramecium quadecaurelia*.

Once the merging for the TGD was complete, we used the *POInT_ances_order* program (26) to infer an ancestral genome order of the resulting 7,669 pillars at the time of the polyploidy event. This program also uses simulated annealing to minimize the number of synteny breaks in the final set of homoeologous genes; we refer to each of these homoeologous loci as a “pillar.” We ran *POInT_ances_order* multiple times until the number of breaks converged. For the paramecium event, we needed to deconvolute the two nested tetraploidies, as described in the next section, after which we used *POInT_ances_order* in a similar fashion.

### Deconvoluting a nested polyploidy

The paramecia genomes studied have undergone two genome duplications since their split with *P. caudatum* (28). Hence, each genome has *four* homoeologous subgenomes present. POInT’s computation of orthologous chromosomal segments scales as *O*(24^2*n*^) for such octoploidy events, making the analysis of more than two such genomes intractable. Instead, we have previously described an approach for deconvoluting the two nested polyploidies to produce a dataset representing only the products of the more recent genome doubling (26).

#### Modeling nested genome duplications

We first seek to phase the four homoeologous regions in each modern *Paramecium* genome into two pairs of homoeologous regions, each descending from one of the two *ancient subgenomes (ASG*) produced by the first tetraploidy (WGD#1). Within each such ASG, there are two *recent subgenomes (RSG)* produced by the second and more recent tetraploidy event (WGD#2). Our approach works by first fitting POInT’s octoploidy (WGQ) model to pairs of polyploid genomes in the set. We construct these pairs such that each genome is present in at least two pairs (Supplemental Table 2). We first fit the WGQ_null_ model with three parameters: a branch length representing the rate of loss from quadruplicated to triplicate loci, and two scaling parameters (δ and σ; Supplemental Figure 1), representing the relative rate of loss from triplicated to duplicated loci (δ), and from duplicated to single-copy loci (σ), respectively. This model serves as a baseline for the test of whether there is sufficient statistical signal in the genomes to confidently identify the two ASGs. Next, to make this test, we fit the WGQ_root_ model to the data. This model explicitly describes the two-step process of octoploidy (Supplemental Figure 1). Rather than having each loci start in a quadruplicated state *Q*, the loci start in a duplicated root state *D*_*1,3*_ (Supplemental Figure 1), representing the genome’s state after WGD#1 but prior to WGD#2. To understand the utility of this root model, we start by considering the case where the second tetraploidy followed immediately after the first. In those circumstances, we would expect to have no duplicate losses prior to WGD#2, meaning that every locus would have passed directly from *D*_*1,3*_ to *Q*. On the other hand, any losses prior to WGD#2 will result in genes that only survive from one of the two ASGs, potentially allowing us to distinguish between them. To detect this lag, we arbitrarily define gene copy 1 to have derived from ASG 1 and copy 3 from ASG 2. If copy 1 is lost prior to WGD#2, then rather than passing to state *Q*, that locus instead passes to state *D*_*3,4*_ because copy 2 was never produced by WGD#2 due to the lack of copy 1. The situation is analogous to the loss of copy 3. The model lets the rate of loss after WGD#1 occur at rate λ, meaning that when λ=0 we fall back to the case of WGD#2 following WGD#1 instantaneously with no intervening losses. We can use a likelihood ratio test (53) with one degree of freedom to test this null hypothesis of λ=0:. When we do so, we find for all pairs of genomes that the WGQ_root_ model is a significant improvement over the WGQ_null_ model (Supplemental Table 2), indicating the WGD#2 occurred after significant gene losses had occurred following WGD#1.

#### Assigning genes to ancient subgenomes

Given that WGQ_root_ is a valid description of the evolution of these genomes, we can use it to probabilistically assign genes to ASGs. Each such assignment is done based on POInT’s conditional orthology probabilities (42), meaning that each set of four genomic locations is given a probability of belonging to ASG 1 or 2. (The WGQ_root_ model is degenerate, so that swapping copies 1&3 with 2&4 gives an identical inference, but this symmetry does not alter our approach).

#### Identifying loci with consistent ASG assignments

We next take the ASG assignments from the pairs in Supplemental Table 2 and seek to merge them. In all cases, those pairwise ASG assignments are required to have orthology confidence *cε*0.95 to be included in our analyses. For a given ancestral locus, we can think of at most four descendant genes present in the modern octoploid paramecia genomes. At such a locus, we start by selecting an arbitrary genome pair G1 and G2 (say *P. decaurelia/P. dodecaurelia)*: we place the genes from each genome assigned to ASG 1 in one list (*L1*) and the genes assigned to ASG 2 in a second list (*L2*). We next proceed to find loci produced by WGD#2 as follows:

□ Find the other genome *P* that *G1* is paired with and add genome pair *G1, P* to the stack of pairs *S*
□ Find the other genome *P* that *G2* is paired with and add genome pair *G2, P* to the stack of pairs *S*
□ While (*S* is non-empty)
  □ Pop a genome pair *Gs, P* from *S*
    □ If (*Gs,P*) has not been processed:
      □ Identify ASG 1 and ASG 2 from *P* based on *Gs* and add the genes from *P* to *L1* and *L2*, respectively
    □ Find the other genome partner *N* of *P* and add *P, N* to *S*
□ If the lists *L1* and *L2* have no common genes
  □ If *L1* has at least one gene from every genome, add those genes as a new locus to our inferred WGD#2
  □ If *L2* has at least one gene from every genome, add those genes as a new locus to our inferred WGD#2

The result of this algorithm is a set of loci duplicated at the most recent WGD event with surviving copies in all the genomes and consistent assignments to ASG 1 and 2 from our model. These loci can then be analyzed with standard tetraploidy models in POInT. In this case, we found 3,925 pillars that we could analyze. We optimized the order of those pillars, resulting in a pillar order with 4,375 breaks.

### Inferring the species tree for POInT analyses

To use POInT to make an estimate of the species trees for taxa sharing these events, we first inferred gene trees for all single-copy genes in our pillar datasets. To do so, we first aligned the protein sequences of all of the genes for each pillar for each event with T-coffee (v13.45) (54). We then found the corresponding nucleotide alignments and inferred the maximum likelihood gene tree with PAUP* (v.4.0, build 168), using the HKY model (55) with empirical base frequencies and site-rate variation modeled with a discrete gamma distribution (56).

The gene trees where none of the taxa had surviving duplicates from the polyploidy (i.e., single-copy genes) were used to rank all the observed topologies in those trees by their frequencies. In other words, we looked at how many pillars supported each observed topology. We then fit the 3-5 most frequent topologies to the observed gene order data using POInT under the WGD_bf_ model (Figure 1) and identified the topology with highest likelihood under that model (Figure 2) (57).

### Evaluating POInT’s models of post-WGD evolution

For each event, we compared four nested models of post-polyploid evolution (Figure 1). In the WGD_null_ model, duplicate losses occur symmetrically from both subgenomes after polyploidy and ohnologs do not become fixed. The WGD_fix_ model adds the possibility of ohnolog fixation, occurring at rate γ, to WGD_null_. The WGD_bias_ model adds a bias towards gene losses from the (inferred) progenitor subgenome #2 (rate ε), meaning that more surviving genes are seen from subgenome #1. Finally, model WGD_bias/fix_ includes both bias in duplicate losses and duplicate fixation. The WGD_null_ model is nested with respect to WGD_fix_ (γ=0) and WGD_bias_ (ε=1), so we used a likelihood ratio test with one degree of freedom to assess whether these two models represented an improvement over WGD_null_. Because WGD_null_, WGD_bias_ and WGD_fix_ are all nested within WGD_bias/fix_, we further conducted likelihood ratio tests comparing these three models to WGD_bias/fix_, with 2 degrees of freedom for the comparison to WGD_null_ and one degree of freedom for the comparisons to WGD_fix_ and WGD_bias_.

### Measuring the clustering of ohnolog losses through time after polyploidy

POInT estimates *conditional probabilities* (43): the probability of transitions between each model state and all other model states at each pillar along each branch of the species tree. Hence, for pillar *x* there is a probability *P*_*U*⟶*S*1_ estimated for the transition *U*⟶*S*_1_ along each branch. Note that this probability could be 0 were the ohnolog pair still present in the taxa in question at pillar *x*. For the models we used, we are interested only in the losses, regardless of the subgenome from which the loss originates. Hence, at each pillar *x* and branch *b* we compute:

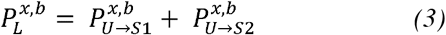

For the yeast dataset, we also included losses from the converging states previously described (43), but the 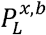 values are entirely equivalent. For each branch, we first calculate *D* the mean (average) absolute difference in 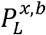 between neighboring pillars:

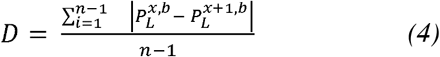

where *n* is the number of pillars. If losses at each pillar are purely independent, we would expect 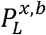 and 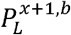 to be no more similar to each other than two randomly selected pillars. Hence, we randomized the pillar order 1,000 times and compute *D*_*rand*_: the difference in loss probability seen between random pillars. The distribution of the *D*_*rand*_ allows us to conduct a statistical test of the null hypothesis that the observed *D* for branch *b* is consistent with independent ohnolog losses along that branch. We define the *P* values for this test to be the number of values of *D*_*rand*_ as small or smaller than *D*.

More usefully, we can use the distribution of the *D*_*rand*_ values to estimate how far from the null distribution *D* is. To do so, we first compute the standard deviation of the distribution of *D*_*rand*_, 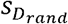:

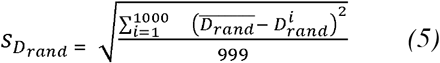

Where 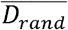 is the mean value of *D*_*rand*_. We define Δ*D* as:

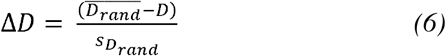

We can also trivially compute *L*^*b*^: the total number of ohnolog losses on branch *b:*

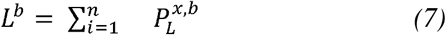

Finally, we can compute *t*^*b*^: the sum of the branch lengths from the root of the tree to the end of branch *b*. In Figure 3, we show the relationship between *t*^*b*^ and *L*^*b*^ and Δ*D*.

### Inference of gene conversion from gene tree topologies

For any pillar with at least one surviving duplicate pair from the WGD, we tested whether the inferred gene tree for that pillar was more compatible with the hypothesis that that pillar evolved under the species tree or had experienced gene conversion (GC) after the WGD. To do so, we defined three hypothesized gene trees: two gene-conversion compatible topologies and a species tree. In the “full gene conversion” case (#1), every surviving ohnolog pair is made to be sister in the tree, corresponding to every duplicate having experienced a gene conversion since its split with its nearest phylogenetic relative (Supplemental Figure 2). In the “early gene conversion case” (tree #2), we hypothesized a conversion event in one or the other of the two basal lineages of the polyploid taxa (Supplemental Figure 2). In other words, we identified the earliest split in the lineages sharing the WGD and assumed that the two lineages produced by that split showed surviving duplicate genes that clustered with their paralogs in that lineage, rather than a gene tree compatible with mirrored copies of the species tree (tree #3; Supplemental Figure 2).

We thus mapped all pillars with a surviving homoeolog pair onto these three alternative topologies, pruning out any lost homoeologs, using POInT’s orthology estimates at a confidence cutoff of *c* ≥ 90%. We computed the Robinson-Foulds (RF) distance (32), implemented in the R package TreeDist (58), between the inferred gene tree for that pillar (from PAUP*) and each of the three hypothesized trees above. We considered potential cases of gene conversion to be those where one or the other of the gene conversion trees had a lower RF distance to the inferred gene tree than did the species tree. We further required that all three trees be topologically distinguishable, meaning that their RF distances to the inferred gene tree were all numerically distinct.

To statistically test for the support for the inference of gene conversion, we next took any pillar meeting the criteria for conversion above and used sequence simulation to assess the evidence for a better fit to the GC tree than the species tree. We thus fit the aligned sequence data to the appropriate GC tree and to the species tree under the HKY model (55) with observed base frequencies. We computed the difference in log-likelihood between the GC and species tree. We then simulated 100 nucleotide alignments from the assumed species tree of the same length as the original alignment. We then fit each simulated alignment to the GC tree and the species tree and computed the difference in log-likelihood between these two trees. We computed the significance of the evidence for gene conversion by asking how often in the simulations the log-likelihood of the GC tree exceeded the log-likelihood of the species tree by as much as was seen for the real data. Our *P* value was then just this number divided by the total of 100 simulations. To be conservative, we only considered the hypothesized gene conversion to have occurred if this *P* value was less than 0.01 and the GC tree had a likelihood of at least 5 log-likelihood units greater than the species tree (see Table 1).

#### Clustering of GC events

To assess if the inferred GC events were clustered in the pillars, we computed the mean distance between those GC events in the list of pillars. We then randomized the pillar numbers and drew the same number of pillars from those randomized pillars 100 times and computed the mean distances for those 100 randomizations, computing a *P* value as the number of times the random pillar orders had a smaller mean distance than seen for the real dataset (Table 1).

### Pairwise estimates of gene conversion

To test for recent gene conversion events within each genome, a triplet-based sequence analysis was used. This approach uses a focal gene (*G*1), its ohnolog (*G*2) and its ortholog (*O*) in the nearest relative in the dataset. These three genes induce a degenerate phylogenetic tree with three branches (Figure 6). Using the codon model of Muse and Gaut/Goldman and Yang, branch-specific values of the synonymous (K_s_) and nonsynonymous (K_a_) divergence are computed by maximum likelihood (i.e., each branch has an independent length and value of the selective constraint parameter K_a_/K_s_) (59–61). We can denote the K_a_ values for *G*1, *G*2 and *O* as *Ka*_1_, *Ka*_2_, and *Ka*_*o*_, respectively. Under the hypothesis of no gene conversion, the common ancestor of *G*1 and *G*2 is at the time of the WGD, which is much older than the speciation event separating *G*1 and *O*. Hence, we expect *Ka*_2_to be the largest of the three K_a_ values. However, if gene conversion is ongoing, the divergence of the two branches leading to the ohnologs will be reduced, potentially yielding the case:

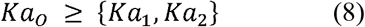

Using POInT (ver. 1.63) (26) we identified ohnolog pairs for which the ortholog of one of the two could be identified with high confidence (*c* ≤ 0.90). The protein sequences of the resulting sequence triplet were aligned with T-coffee (ver. 13.45) (54) and the nucleotide alignment deduced. The nucleotide alignment was fit to the unrooted tree of Figure 6A and any triplets satisfying Equation 8 were then tested for statistical significance with a likelihood ratio test (53). Specifically, we defined the null and alternative models *H*_0_ and *H*_*a*_ as:

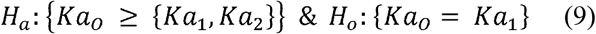

We assumed that the log of the difference likelihood between the two models 2(*ln H*_0_ − *ln H*_*a*_) followed a chi-squared distribution with 1 degree of freedom, as *H*_*a*_ has one more parameter (value of K_a_) than does *H*_0_. The pairwise synonymous divergence of the ohnolog pair was computed with the same codon model described above. If an ohnolog pair has low K_s_ value, this observation further suggests recent gene conversion (62).

## Acknowledgements

This work was supported by U.S. National Science Foundation grant NSF-DEB-2241312.

## Data availability

The POInT and like_tri_test packages are available from GitHub (https://github.com/gconant0/POInT and https://github.com/gconant0/like_tri_test, respectively). Gene order data, POInT models, visualizations, and pillar sequences are all available from the POInT_browse_ web portal (https://wgd.statgen.ncsu.edu).

## Abbreviations

WGD: whole genome duplication,
DCS: double-conserved synteny,
GC: gene conversion,
TGD: teleost whole genome duplication

## Notes

### Competing Interest Statement

The authors have declared no competing interest.

### Summary of Updates

I have added the Supplemental Files. The manuscript and all information the document contains is the same as the previous submission.

