## Supplemental Tables and Figures for "Evidence for post-allopolyploidy genetic exchanges between duplicated regions in three ancient polyploidies"

**Supplementary Information**

SUPPLEMENTAL TABLES:

**1:**

|  | **Duplicates: Focal and Paralog** \| **Ortholog** \| **# GC Pillars** |
| --- | --- |
| **A** | *Paramecium_sexaurelia* \| *Paramecium_sonneborni* \| 68; *Paramecium_sonneborni* \| *Paramecium_sexaurelia* \| 217; *Paramecium_pentaurelia* \| *Paramecium_primaurelia* \| 13; *Paramecium_tetraurelia* \| *Paramecium_octaurelia* \| 17; *Paramecium_octaurelia* \| *Paramecium_tetraurelia* \| 9; *Paramecium_biaurelia* \| *Paramecium_tetraurelia* \| 88; *Paramecium_tetraurelia* \| *Paramecium_biaurelia* \| 64; *Paramecium_tredecaurela* \| *Paramecium_primaurelia* \| 12; ***Paramecium_primaurelia*** \| ***Paramecium_tredecaurela*** \| **101**; *Paramecium_dodecaurelia* \| *Paramecium_decaurelia* \| 6; *Paramecium_decaurelia* \| *Paramecium_dodecaurelia* \| 4 |
| **B** | *Anabas_testudineus* \| *Betta_splendens* \| 29; *Astyanax_mexicanus* \| *Danio_rerio* \| 38; *Betta_splendens* \| *Anabas_testudineus* \| 22; *Danio_rerio* \| *Astyanax_mexicanus* \| 93; *Mastacembelus_armatus* \| *Anabas_testudineus* \| 14; ***Maylandia_zebra*** \| ***Xiphophorus_maculatus*** \| **60**; *Oryzias_latipes* \| *Xiphophorus_maculatus* \| 11; *Parambassis_ranga* \| *Xiphophorus_maculatus* \| 44; *Sparus_aurata* \| *Anabas_testudineus* \| 13; *Sphaeramia_orbicularis* \| *Xiphophorus_maculatus* \| 47; *Xiphophorus_maculatus* \| *Oryzias_latipes* \| 36 |
| **C** | *S_cerevisiae* \| *S_uvarum* \| 67; *S_uvarum* \| *S_cerevisiae* \| 64; *S_kudriavzevii* \| *S_cerevisiae* \| 36; *C_glabrata* \| *S_cerevisiae* \| 66; *K_africana* \| *K_naganishii* \| 172; *K_naganishii* \| *K_africana* \| 107; ***N_castellii*** \| ***N_dairenensis*** \| **104**; *N_dairenensis* \| *N_castellii* \| 89; *V_polyspora* \| *T_phaffii* \| 142; *V_polyspora* \| *T_blattae* \| 198; *T_phaffii* \| *V_polyspora* \| 63; *T_blattae* \| *V_polyspora* \| 59 |

**Supplemental Table 1**: Indicates the use of the pairwise estimates of gene conversion using the triplet test described in the Methods section, where two WGD produced duplicate genes (focal and paralog) are found and tested against an ortholog gene to find areas where gene conversion (GC) is possible. All estimates include significant and non-significant values. For completeness, every pair was tested twice with each possible gene as the focal/paralog and ortholog. If any pairs are missing from the data, that indicates the topology of those pairs are incompatible. The pairs in bold have corresponding plots in Fig.5 of the main manuscript.

**2:**

| **Genome 1^a^** | **Genome 2 ^a^** | **WGQ_null_ lnL^b^** | **WGQ_root_ lnL^c^** | ***P* value^d^** |
| --- | --- | --- | --- | --- |
| *P. decaurelia* | *P. dodecaurelia* | -41012.43 | -39379.55 | ${10}^{-10}$ |
| *P. biaurelia* | *P. tredecaurelia* | -48474.50 | -45339.84 | ${10}^{-10}$ |
| *P. pentaurelia* | *P. primaurelia* | -41966.67 | -39497.86 | ${10}^{-10}$ |
| *P. sexaurelia* | *P. sonneborni* | -50753.20 | -46270.38 | ${10}^{-10}$ |
| *P. tetraurelia* | *P. octaurelia* | -44811.78 | -41626.14 | ${10}^{-10}$ |
| *P. decaurelia* | *P. pentaurelia* | -40036.94 | -37655.80 | ${10}^{-10}$ |
| *P. dodecaurelia* | *P. tetraurelia* | -45247.65 | -42481.16 | ${10}^{-10}$ |
| *P. primaurelia* | *P. sonneborni* | -50886.09 | -46519.89 | ${10}^{-10}$ |
| *P. sexaurelia* | *P. tredecaurelia* | -50408.94 | -46630.24 | ${10}^{-10}$ |
| *P. biaurelia* | *P. octaurelia* | -46448.1512 | -43155.94 | ${10}^{-10}$ |

**Supplemental Table 2:**  Indicates results of octoploid models fit to pairs of *Paramecium* genomes. **a)** Two *Paramecium* genomes that are octoploid relative to the outgroup *Paramecium caudatum.* **b)** Log-likelihood of the quadruple-conserved synteny data for these two genomes relative to *P. caudatum* under a model of instantaneous back-to-back tetraploidy (see Supplemental Figure 1). **c)** Log-likelihood of the quadruple-conserved synteny data for these two genomes relative to *P. caudatum* under a model where there was a lag between the first and second tetraploidies (Supplemental Figure 1). **d)** *P*-value of the hypothesis test of no significant improvement of the WGQ_root_ model over the WGQ_null_ model. Chi-square test with 1 degree of freedom.

SUPPLEMENTAL FIGURES:

**1:**

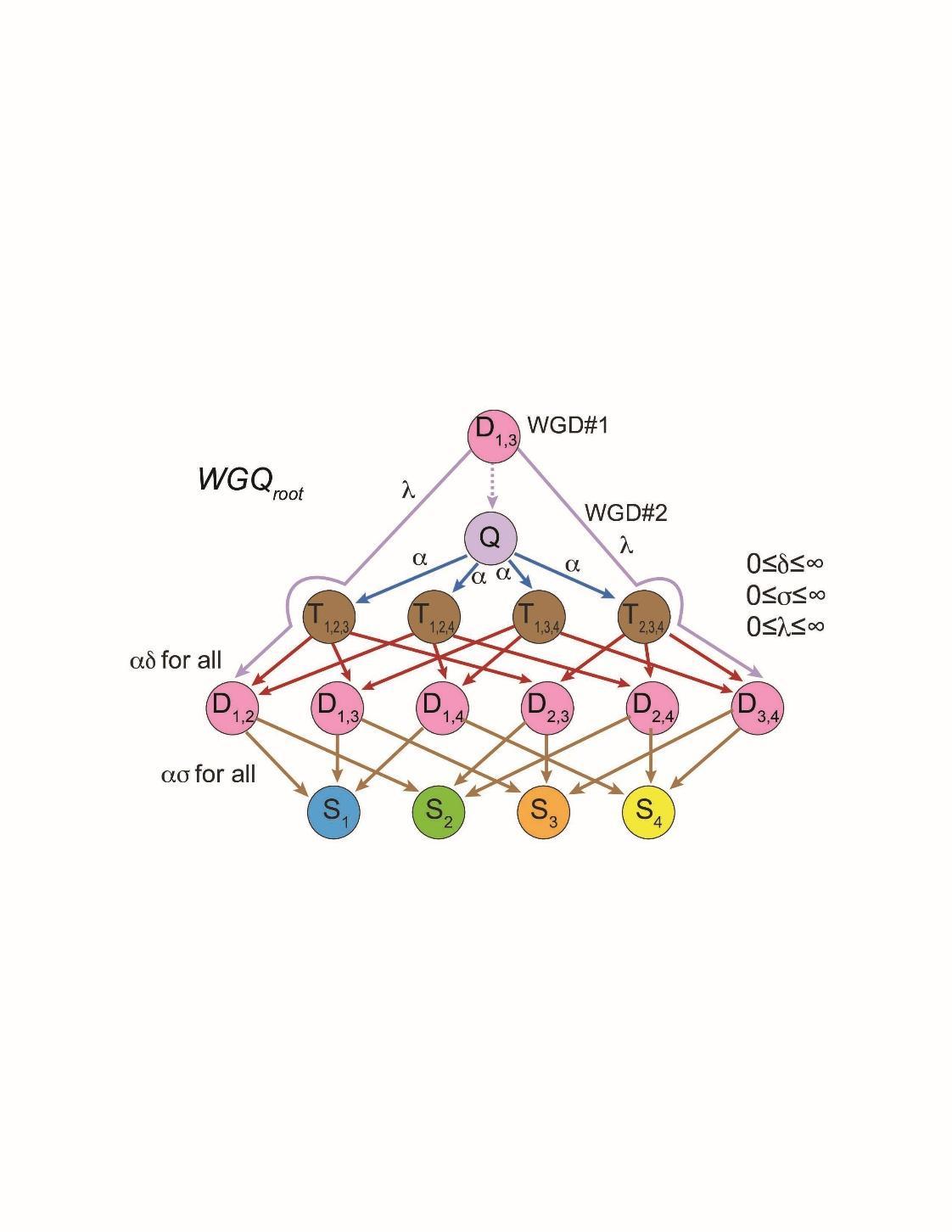

**Supplemental Figure 1:** A model for the resolution of two nested tetraploidy events, as seen in the ciliates. In the base model (*WGQ_null_*, not pictured), all homoeologous loci become quadruplicated instantaneously (state *Q*): losses then proceed through loss of gene copies through the triplicated (*T_x,y,z_*), duplicated (*D_x,y_*) and single copy (*S_x_*) states. If we allow for a gap in time between the two tetraploidies (*WGQ_root_*, pictured), we can assume that all loci start in state *D_1,3_*, corresponding to the first tetraploidy. Prior to the second event, a loss of copy #1 means that the second tetraploidy will only duplicate copy #3, producing a locus in state *D_3,4_* (and symmetrically with copy #3 and state *D_1,2_*). The degree of duplicate loss after the first tetraploidy is modeled with the λ parameter: when λ=0, the *WGQ_root_* falls back to the *WGQ_null_* model. All parameters (λ, as well as the triplication and duplication loss rates, δ and σ) are estimated from the synteny-block data by POInT with maximum likelihood.

**2:**

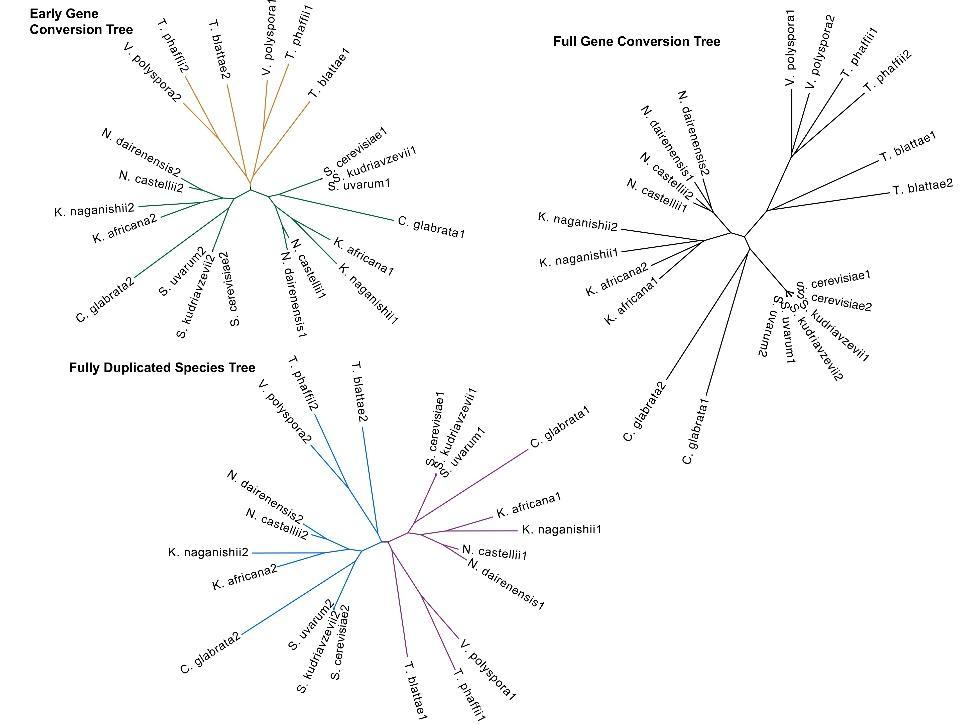

**Supplemental Figure 2:** Examples of the expected gene tree topologies from duplicated loci under three different hypotheses. At the bottom is the expectation in the case of no gene conversion: namely two mirrored copies of the species tree (shown unrooted with blue and purple branches). In the case of complete recent gene conversion (right), each homoeolog is sister to its paralog in the same genome. Finally, in the early gene conversion case (upper left), the first split between the species is assumed to be followed by a gene conversion in one or both lineages, meaning that these two lineages have homoeologous groups that are sister to each other, rather than the mirrored copies of the species seen at bottom.

**3:**

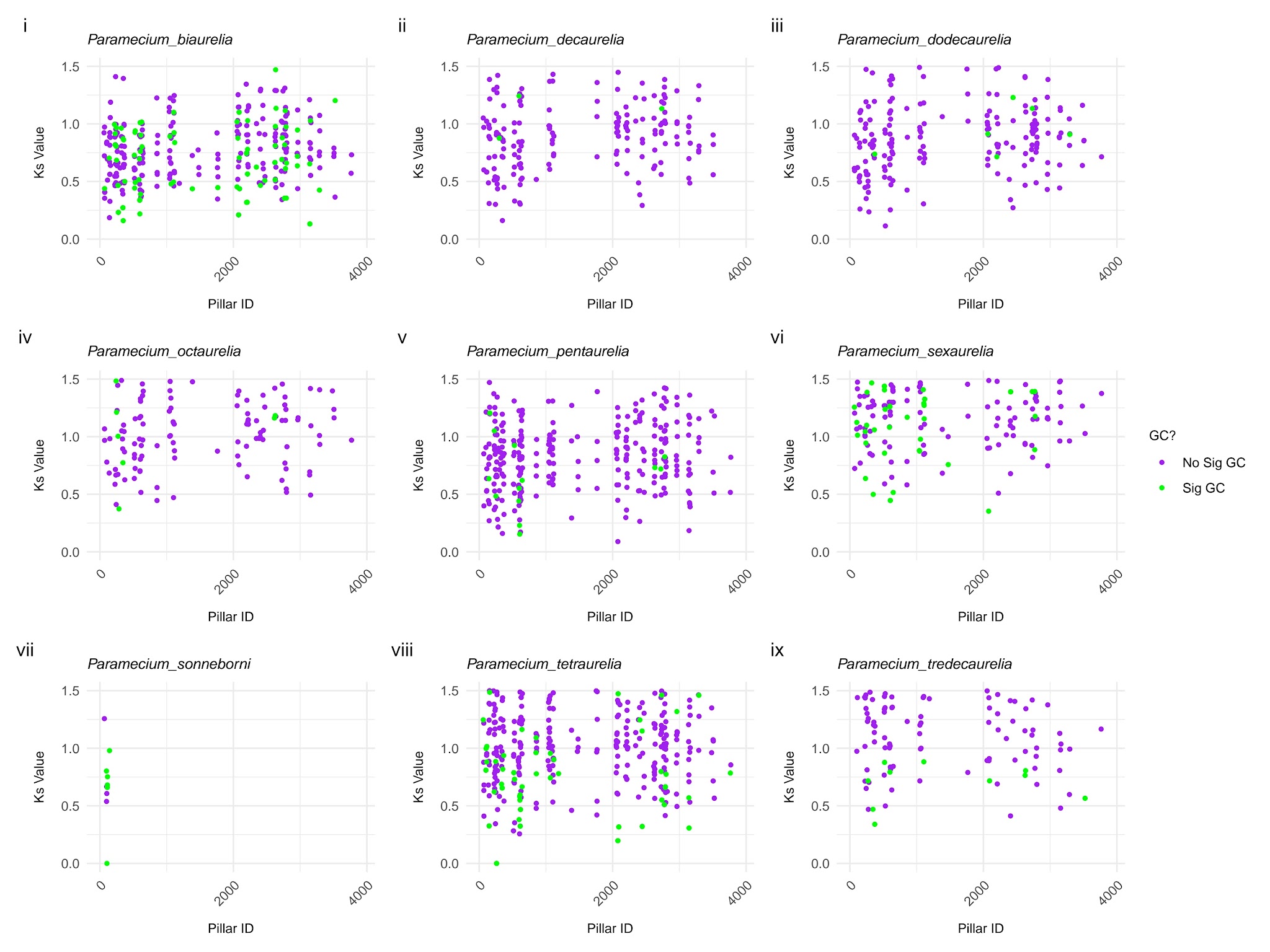
**Supplemental Figure 3:** Pairs of ciliate paramecium variants mentioned in Supplemental Table 1A. Each plot indicates statistically significant gene conversion cases in green, and all other ohnolog pairs are shown in purple. Statistically significant cases of gene conversion are indicated in green $(P<0.05)$; all other ohnolog pairs are shown in purple. The *x* axis shows the position in ancestral genome order; on *y* is the synonymous divergence between the ohnolog pair ($K_{s}$, or the number of synonymous substitutions per synonymous site, *Methods*); ohnolog pairs with K_s_ greater than 1.5 are omitted. Only high confidence (90%) pillars taken from POInT of ohnolog pairs were plotted.

**4:**

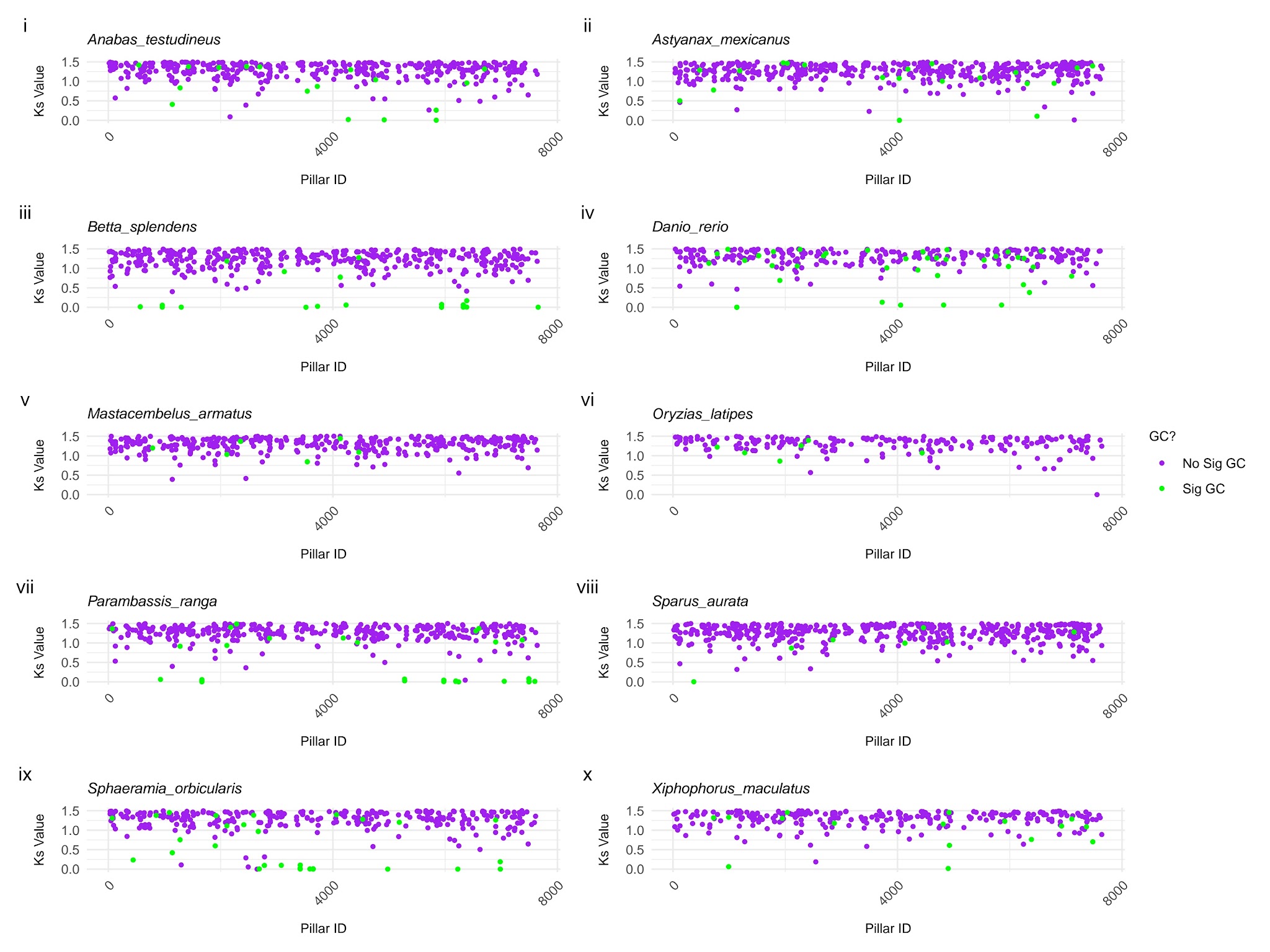
**Supplemental Figure 4:** Remaining pairs of TGD (Bony Fish) Variants mentioned in Supplemental Table 1B. Each plot indicates statistically significant gene conversion cases in green, and all other ohnolog pairs are shown in purple. Statistically significant cases of gene conversion are indicated in green $(P<0.05)$; all other ohnolog pairs are shown in purple. The *x* axis shows the position in ancestral genome order; on *y* is the synonymous divergence between the ohnolog pair ($K_{s}$, or the number of synonymous substitutions per synonymous site, *Methods*); ohnolog pairs with K_s_ greater than 1.5 are omitted. Only high confidence (90%) pillars taken from POInT of ohnolog pairs were plotted.

**5:**

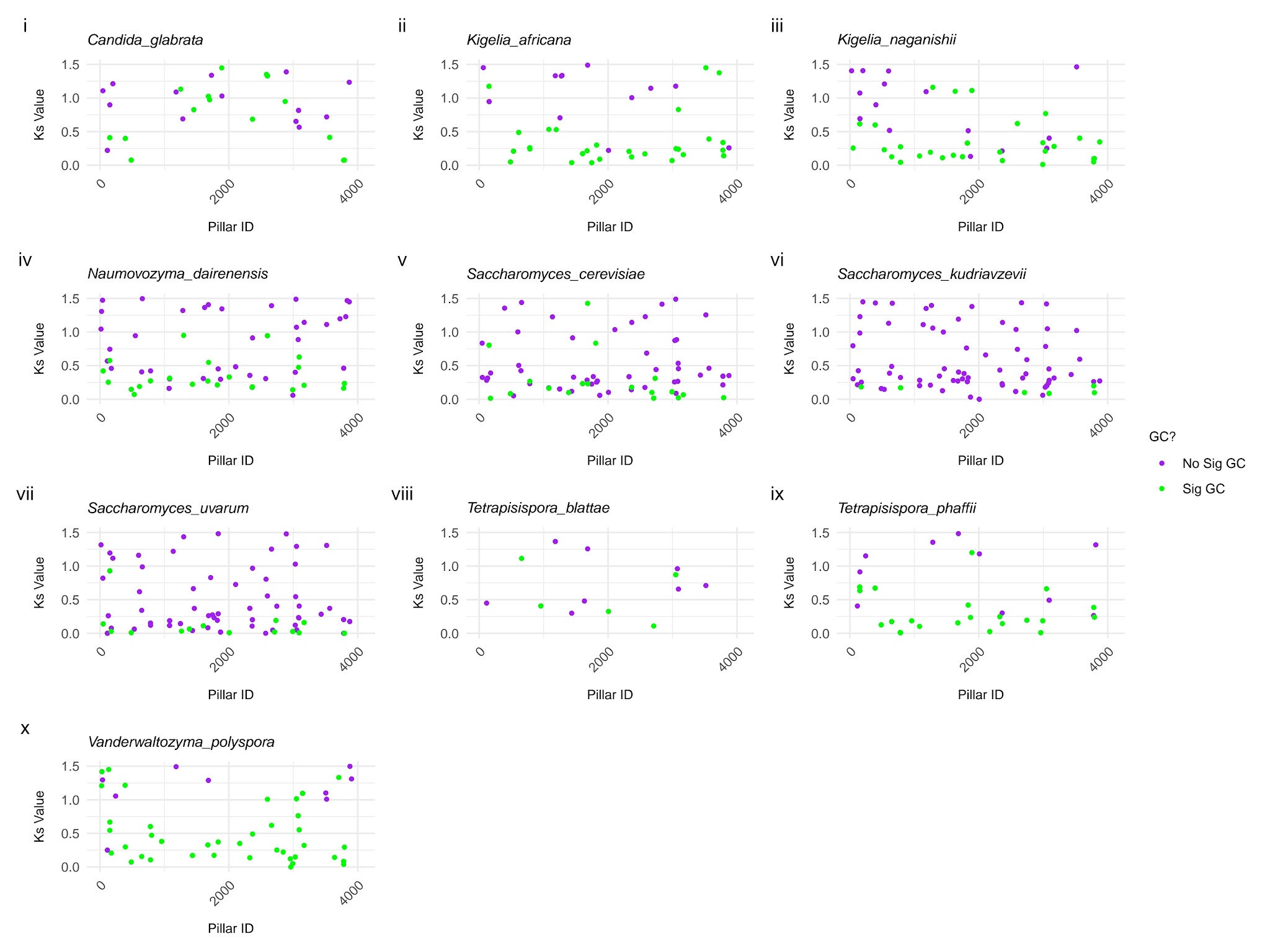
**Supplemental Figure 5:** Remaining pairs of Baker’s Yeast Variants mentioned in Supplemental Table 1C. Each plot indicates statistically significant gene conversion cases in green, and all other ohnolog pairs are shown in purple. Statistically significant cases of gene conversion are indicated in green $(P<0.05)$; all other ohnolog pairs are shown in purple. The *x* axis shows the position in ancestral genome order; on *y* is the synonymous divergence between the ohnolog pair ($K_{s}$, or the number of synonymous substitutions per synonymous site, *Methods*); ohnolog pairs with K_s_ greater than 1.5 are omitted. Only high confidence (90%) pillars taken from POInT of ohnolog pairs were plotted.

**6:**

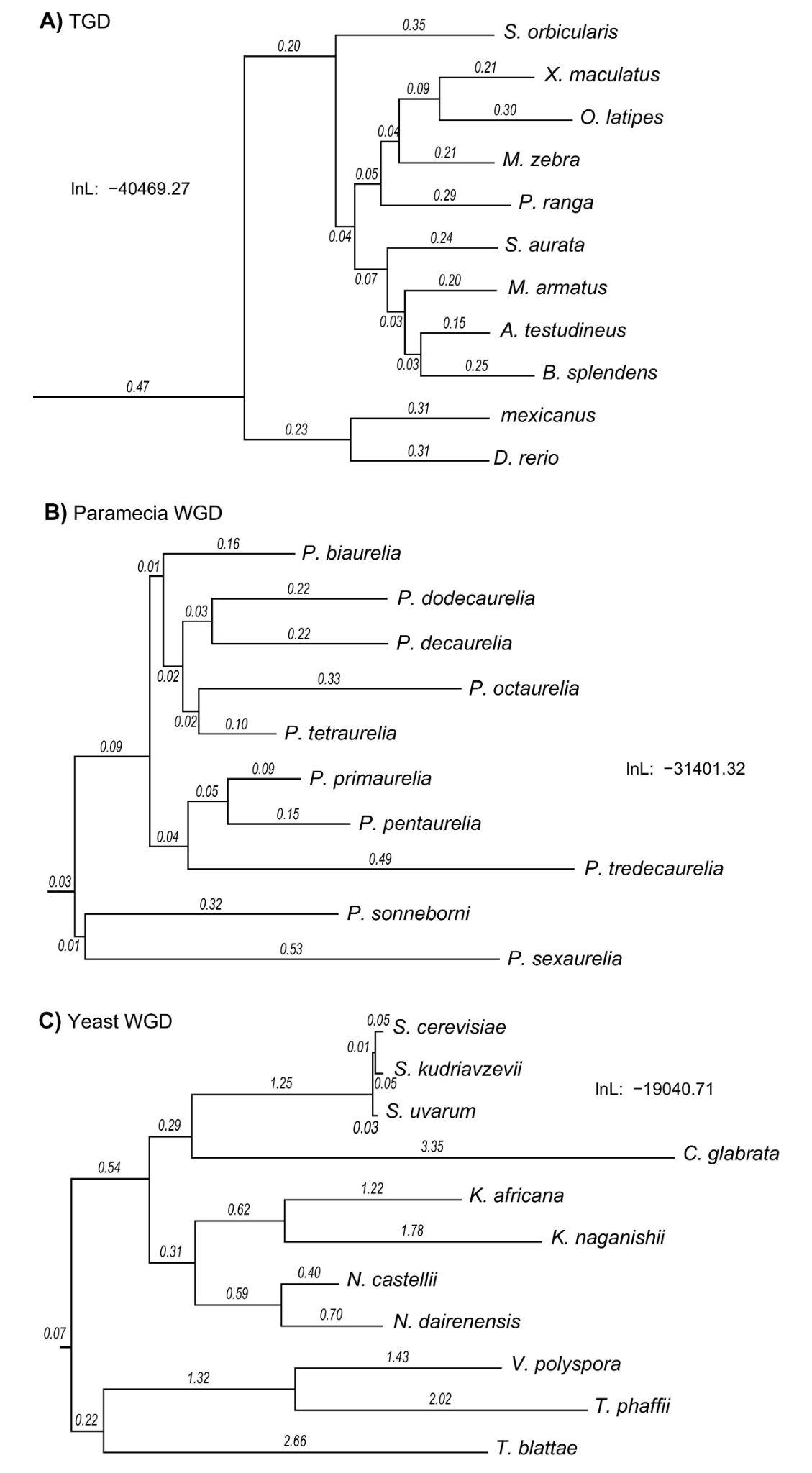

**Supplemental Figure 6:** Inferred species tree topologies from POInT for each event with scaled branch lengths. Branch lengths represent the inferred rate of ohnolog loss along a branch. **A)** The TGD under the WGD_bf_ model. **B)** The paramecia WGD event under the WGD_f_ model. **C)** The yeast WGD under the WGD_cf_ model (includes converging losses, see Emery et al., cited in the main manuscript.)
